# *Keetia gordonii* sp. nov. (Rubiaceae - Vanguerieae) a new species of threatened forest liana from the littoral forest of Gabon

**DOI:** 10.1101/2024.04.18.590134

**Authors:** Martin Cheek, Pulchérie Bissiengou

## Abstract

*Keetia gordonii* sp. nov. (Rubiaceae - Vanguerieae) a new species of forest liana from the littoral forest of Gabon is formally described and illustrated as the first endemic species of the genus from that country. On current evidence, the species appears to have three locations and is threatened by forest clearance. It is provisionally assessed using the IUCN 2012 standard as Endangered (EN B2a,b(i-iv). The new species is extremely distinctive within the genus, showing several character states previously unrecorded in *Keetia. Keetia gordonii* is currently unique in its genus for the massively thick, coriaceous leaf blades seen in the fruiting axillary branches (in this respect resembling a *Psydrax* Gaertn., vs papery or rarely thinly coriaceous in other *Keetia* species), and also for the globose, smooth pyrenes (vs ovoid, colliculose in other species), in which the lid crest is so vestigial that it is almost imperceptible (vs lid crest conspicuous in other species). The tanniferous seed endosperm shows a new character state for the genus being present in a continuous, solid layer in the outer part of the seed, rather than being in radial bands or diffuse as in other species of the genus. However, there is no doubt that this taxon is best placed in *Keetia* as opposed to *Psydrax* due to the disc concealed deep inside the calyx tube (vs exposed), the presence of a pyrene cap (vs none) and the stipules that lack a well-developed keel (vs keel present). Further, the presence of tanniniferous seed endosperm is not recorded in any other genus of the tribe. *Keetia gordonii* is currently assigned to the *K. hispida* species group of Guineo-Congolian Africa.

## Introduction

*Keetia* E. Phillips (Rubiaceae Vanguerieae) was segregated from *Canthium* Lam. by Bridson (1985, 1986). Restricted to sub-Saharan continental Africa (absent from Madagascar and the Mascarene Islands), and extending from Senegal and Guinea in West Africa (Gosline *et al*. 2023a; 2023b) to Sudan in the North and East (Darbyshire *et al*. 2015) also Ethiopia, and S. Africa in the South (Bridson 1986), this genus of about 41 accepted species (Cheek & Onana 2024) are distinguished from similar Canthioid genera in west Africa by their pyrenes with a fully or partly-defined lid-like area around a central crest and endosperm with tanniniferous areas (Bridson 1986). *Keetia* species are mainly climbers of forest (rarely shrubs of wooded grassland or small trees). In a phylogenetic analysis of the tribe based on morphology, nuclear ribosomal ITS and chloroplast *trnT-F* sequences, Lantz & Bremer (2004), found that based on a sample of four species, *Keetia* was monophyletic and sister to *Afrocanthium* (Bridson) Lantz & B. Bremer with strong support. Highest species diversity of *Keetia* is found in Cameroon and Tanzania, both of which have about 15 taxa (Onana 2011; POWO, continuously updated). In contrast, in neighbouring Gabon there are only 10 species, although more than two thirds of specimens recorded in that country remain unidentified to species (Sosef *et al*. 2006), indicating the current incomplete knowledge of the genus in Gabon. Several *Keetia* species are point endemics, or rare national endemics, and have been prioritized for conservation (e.g. Onana & Cheek 2011; Couch *et al*. 2019; Murphy *et al*. 2023; Darbyshire *et al*. 2023) and one threatened species, *Keetia susu* Cheek has a dedicated conservation action plan (Couch *et al*. 2022)

Bridson’s (1986) account of *Keetia* was preparatory to treatments of the Vanguerieae for the Flora of Tropical East Africa (Bridson & Verdcourt 1991) and Flora Zambesiaca (Bridson 1998). Pressed to deliver these, she stated that she could not dedicate sufficient time to a comprehensive revision of the species of *Keetia* outside these areas: “full revision of *Keetia* for the whole of Africa was not possible because the large number of taxa involved in West Africa, the Congo basin and Angola and the complex nature of some species would have caused an unacceptable delay in completion of some of the above Floras” (Bridson 1986). Further “A large number of new species remain to be described.” Several of these new species were indicated by Bridson (1986), and other new species by her arrangement of specimens in folders that she annotated in the Kew Herbarium. One of these species was later taken up and published by Jongkind (2002) as *Keetia bridsoniae* Jongkind. In the same paper, Jongkind discovered and published *Keetia obovata* Jongkind based on material not seen by Bridson. Based mainly on new material, additional new species of *Keetia* have been published by Bridson & Robbrecht (1993), Bridson (1994), Cheek (2006), Lachenaud *et al*. (2017), Cheek *et al*. (2018a), Cheek & Bridson (2019) and Cheek & Onana (2024), and several other taxa that fit no other species, (e.g. Cheek et al. 2004; 2011)) remain to be described.

In this paper we continue the project towards an updated taxonomic revision of *Keetia* by describing a further new species that was first indicated as such by Bridson (*McPherson* 17011 (K) annotated by Bridson as “species not matched, July 2000”).

The new species, *K. gordonii* Cheek, is remarkable for being the first accepted species of the genus to be described from Gabon (all other recorded Gabonese *Keetia* species were first discovered in other countries), and also to be, on current evidence, the first and so far, only *Keetia* species published as endemic to that country. *Keetia gordonii* is currently unique in its genus for its massively thick, coriaceous leaf blades (in this respect resembling a *Psydrax* Gaertn., vs papery in most other *Keetia* species), and also for the globose, smooth pyrenes (vs ovoid, colliculose in other species), in which the lid crest is so vestigial that it is almost imperceptible (vs lid crest conspicuous in other species). However, there is no doubt that this taxon is best placed in *Keetia* as opposed to *Psydrax* due to the highly tanniniferous seed endosperm (which shows a unique state in the genus, being continuous, see below), the disc concealed deep inside the calyx tube, the presence of a pyrene cap and the stipules that lack a well-developed keel. The species probably belongs to the *K. hispida* (Benth.)Bridson group (see comments below).

New plant species to science such as *Keetia gordonii* continue to be published steadily from Gabon. Recently published new species range from trees and shrubs (Barberá et al. 2023; Couvreur et al. 2022; Dagallier et al. 2023; Lachenaud & Bidault 2022; Quintanar et al. 2023) to epiphytic herbs (Farminhão et al. 2023), epiphytic parasitic shrubs (Lachenaud 2023) to lianas (Jongkind & Lachenaud 2022).

## Materials and methods

The specimen was collected using the patrol method (e.g. Cheek & Cable 1997). Herbarium material was examined with a Leica Wild M8 dissecting binocular microscope fitted with an eyepiece graticule measuring in units of 0.025 mm at maximum magnification. The drawing was made with the same equipment with a Leica 308700 camera lucida attachment. Pyrenes were prepared by boiling selected ripe fruits for several minutes in water until the flesh softened and could be removed. Finally, a toothbrush was used to clean the pyrene to expose the surface sculpture. Names of species and authors follow IPNI (continuously updated). Identification and naming follow Cheek in Davies *et al*. (2023). Specimens were searched for in the following herbaria: BM, BR, K, LBV, P, WAG, YA either in person or using images available online.

The format of the description follows those in other papers describing new species of *Keetia*, e.g. Cheek & Bridson (2019), Cheek & Onana (2024). Terminology follows Beentje & Cheek (2013). Herbarium codes follow Index Herbariorum (Thiers, continuously updated). Nomenclature follows Turland *et al*. (2018). All specimens seen are indicated as “!”. The extent of occurrence was calculated using Geocat (Bachmann et al. 2011) and the conservation assessment follows the IUCN (2012) standard.

## Results

Among the known *Keetia* species recorded from Gabon, all of which are relatively widespread, the new species is most likely to be confused with the common *Keetia gueinzii* (Sond.) Bridson, which is also the most widespread species of the genus (Cameroon to S. Africa, Bridson 1986). The two species share more or less dark reddish brown, patent, conspicuously hairy stems, oblong-lanceolate leaf blades which are usually symmetrically cordate at the base, and with a similar number of secondary nerves. However, the new species also has similarities with the *K. hispida* group (of which it seems to be part, see notes on this group below) especially one element of the *K. hispida* complex itself, ‘setosum’ (Cheek et al. 2004: 375), which also often have cordate, hairy leaf blades of similarly large sizes, and sometimes hairy stems. Other formally published species of the genus in Gabon have more or less glabrous stems and/or much smaller or differently shaped leaf blades. Diagnostic characters separating the new species from *K. hispida s*.*l. ‘setosum’* and *Keetia gueinzii* are set out in Table 1.

**Table 1.** Diagnostic characters separating *Keetia gordonii sp. nov*. from *Keetia gueinzii* and *K. hispida s*.*l*.

|  | *Keetia gueinzii* | *Keetia gordonii*<br>sp.nov. | *K. hispida* s.l.<br>‘*setosum*’ |
| --- | --- | --- | --- |
| Tertiary nerve visibility in leaf blades (abaxial surface) | Conspicuous | Not visible | Conspicuous |
| Leaf blade indumentum (aaaxial surface) | Densely appressed hairy | Very sparsely and inconspicuously erect hairy | Moderately densely appressed hairy |
| Leaf blade texture | Chartaceous | Coriaceous | Chartaceous |
| Stipule shape /<br>Length: breadth | Narrowly<br>triangular to<br>lanceolate / <2:1 | Broadly triangular/<br>c. 1:1 | Narrowly<br>triangular / <2:1 |
| Inflorescence<br>/Infructescence branching | First to third order<br>branches<br>conspicuous | Branches absent or<br>inconspicuous or,<br>if present, first<br>order branches<br>short. | First to third order<br>branches<br>conspicuous |
| Fruiting calyx tube: teeth<br>length | Calyx tube < teeth | Calyx tube > teeth | Calyx tube > teeth |
| Nodes of primary stems | Not inflated or<br>inhabited by ants | Not inflated or<br>inhabited by ants | Inflated, and<br>inhabited by ants |

### Taxonomic treatment

#### Keetia gordonii *Cheek* sp. nov. Figs. 1.& 2

Type: Gabon, Ogooue-Maritime, 32 road-km N of Igotchi-Mouenda, Bakker forestry concession; lowland forest “woody vine; fruit green”. 02° 41’ S, 010° 30’ E, 250 m, fr. 16 May 1997, *McPherson* 17011 (holotype K!, barcode K000593367; isotypes LBV!, MO (Cat No. 1188793), UPS (Cat No. V-142868)).

*Evergreen climber*, climbing to several metres high. *Primary stems* terete, c.0.6 cm of diam., densely mostly reddish brown hairy. *Secondary shoots* (brachyblasts, plagiotropic or spur shoots) leafy, at least c. 30 cm long, with at least 4 internodes, internodes solid, terete, 4.5 − 7 x c. 0.4 cm, (Fig. 1A), at fruiting stage densely dark purplish brown tomentose, hairs dense, appressed to patent, completely covering the surface, 0.2 − 2 mm long, wiry, simple, sometimes wrapping around each other, hairs-hairpin-shaped, kinked or rarely straight, the shorter hairs almost always sinuous. *Leaves* dimorphic, those of primary axis opposite and equal at the nodes, orbicular to ovate, c. 8.3 x 6.4 cm, lateral nerves c. 5 pairs on each side of the midrib, digitally 5-nerved at base, apex acuminate, base broadly cordate; leaves of secondary shoots opposite and equal at each node, distichous, not leathery, blades drying black on upper surface, orange-brown on lower surface, oblong-elliptic, rarely ovate-elliptic or obovate-elliptic, (11.5 −) 15.5− 18.5 x 7 − 8 (− 10) cm, acumen acute triangular (0.8 −)1.0 (− 1.3) x 0.5 − 1 cm; base symmetrical or slightly asymmetric, cordate, sinus angle 90 − 110°, 0.1 − 0.9 x 0.8 − 12 cm, margin slightly thickened, with long hairs; *bacterial nodules* absent (Fig. 1C); midrib raised above and below, densely red brown tomentose, hairs as stem, but shorter, secondary nerves raised and tomentose abaxially, glabrescent and sunken adaxially, domatia absent; secondary nerves 7 −− 9 on each side of the midrib, arising from the midrib, at 60− 70°, straight, but near the margin abruptly looping up and uniting with the nerve above, weakly in the proximal nerves, otherwise strongly, the nerve junctions 3 − 11 mm from the margin; tertiary nerves subscalariform, 3 − 4 connecting the more proximal secondaries, curved or angled; areolae oblong, c. 20 x 6 − 7 mm, quaternary nerves inconspicuous; abaxial surface subscabrid, hairs discernible with the naked eye. hairs moderately dense, c. 40 % cover along the midrib and secondary nerves, simple, pale bronze-coloured, 0.4 − 0.7 mm long, substrigose, straight; intersecondary areas (abaxial surface) sparsely hairy, hairs c. 2 per mm^2^, c. 5 % cover, hairs appressed, curved (Fig. 1B); adaxial surface very sparsely and inconspicuously hairy, hairs erect, curved, c. 0.75−1 mm long, densest on the nerves, surface with sunken golden discs c. 30 per mm^2^, each 0.01 mm diam. (these may be bases of fallen hairs. *Petioles* terete to slightly plano-convex in transverse section, (1 −) 1.2 − 1.5 x 0.2 cm, indumentum densely black hair, hairs as stems. *Stipules* free, wider than the stem, concave, the upper part displaced laterally and curving around the stem, in life green, drying papery, brown, moderately persistent (persisting at third to fourth nodes from apex at fruiting stage), triangular, those of the primary stem c. 3 x 3.3 cm, those of the secondary stem 1.5 − 2.5(− 3) x 1 − 2 cm (at apical bud slightly smaller), tapering gradually to the linear, c. 5 mm long often twisted, apex, awn absent, abaxial (outer) surface (Fig. 1C), hairy, midrib raised, indumentum as midrib, blade densely hairy, hairs bimodal: long, strigose hairs c. 0.4 mm long, sparse; interspersed with dense 0.08 − 0.16 mm long, appressed hairs (Fig. 1D); inner surface glabrous apart from a dense line of hairs and colleters at the base (exposed and remaining attached to the stem when the stipule falls), hairs erect, brown, simple c. 2 mm long mixed with a few sparse colleters; colleters black, botuliform, 0.4 −0.8 x 0.2 mm, apex rounded. *Inflorescences* not seen at anthesis (only old flowers), axillary on spur (plagiotropic) branches, held above the stem, in opposite axils in c. 2 successive nodes, beginning 1 node below stem apex; subcapitate (inflorescence branches highly contracted and inconspicuous, up to 5 mm long), 5 − 9-flowered. Peduncle 10 − 15 x 1 − 1.5 mm, with bract at the apex, bract elliptic 0.5− 1.2 x 0.3 mm indumentum and surface as stipules. Pedicels 2.5 − 3.5 mm long, 0.5 − 1 mm wide, widest at apex, with moderately dense spreading to ascending simple hairs (0.5 −) 0.7 − 1 mm long. Calyx-hypanthium subcylindrical 5 − 5.5 x 2–2.25 mm, calyx tube cylindrical, 1.8 −3 x 2.25 mm long; teeth 5, broadly triangular, c. 1.5 x 2 mm, outer surface densely hairy, with hairs as pedicel (Fig. 1E), inner surface glabrous. Disc glabrous, convex, c. 2 mm diam., 0.5 mm long, the style base sunken. *Infructescences* (Fig. 1A), subcapitate 3 − 4 x 3 − 5 cm, (2−) 5 − 8-fruited, pedicels (accrescent) terete, with 6–7 longitudinal ridges, (5 −)7.5 x 2.5 mm, with moderately dense ascending, curved, simple hairs (0.5 −) 1 − 1.75 mm long. *Fruit* green (potentially immature?), fleshy, fruit wall c. 1. 5 mm thick, inconspicuously didymous, the two carpels united along their length, divided only by a very shallow, barely discernible longitudinal groove on each side (Fig. 1F), in side view suborbicular (excluding calyx), (11−) 12 − 13(− 14) x 11 − 14 x 10 mm, apex with persistent, accrescent, shortly cylindric fleshy calyx c. 2 − 3 x 5 mm; teeth persistent, sometimes completely concealed by dense forward pointing bristle hairs 2− 2.5 mm long (Fig. 1F), disc inconspicuous; base of fruit slightly cordate or rounded; surface smooth, sparsely hairy, 5− 10% cover, with a mixture of patent 2 mm long bristle hairs and crisped 0.4−0.6 mm finer hairs. 1-seeded fruits (by abortion, the minority, 1/3 of all ripe fruit), ellipsoid, symmetric, 12− 14 x 10 x 10 mm. Pyrene pale brown, woody, 0.5− 0.6 mm thick, ellipsoid, c.11 x 9 x 9 mm, the surface smooth apart from the longitudinal narrowly ellipsoid, raised, boss like point of attachment on the ventral face (Fig. 1I). Lid apical, cap-like, orbicular, c. 2.5 x 6.25 x 6 mm, angled c. 20 degrees towards the ventral face, crest (keel) vestigial, barely perceptible. Fig. 1H-J. *Seed* ellipsoid c. 8 x 7 x 5 mm, surface black, convoluted like a brain, epidermal cells finely reticulate; embryo (transverse section) nearly the full width of the seed (c. 7 mm) surrounded in the centre of the seed by a small white powdery area; tanniniferous area not in rays, or dispersed, but a solid, hard, glossy black area forming the marginal 1.5 − 2 mm of the seed (Fig. 2).

**Fig. 1.**
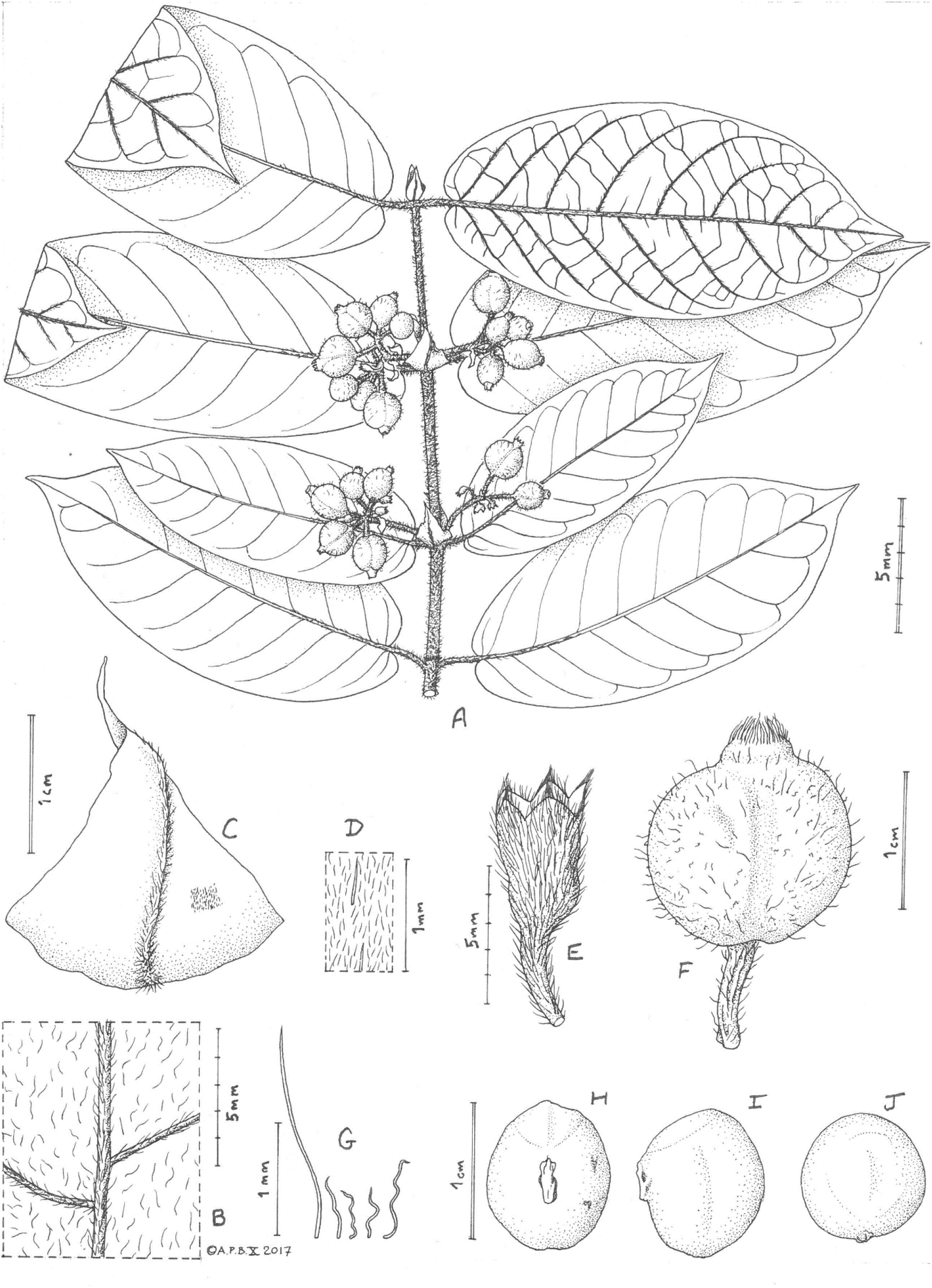
*Keetia gordonii* **A** habit, fruiting secondary axis; **B** leaf blade, abaxial surface showing midrib and secondary nerves; **C** stipule, abaxial surface; **D** detail of C showing indumentum; **E** flower showing calyx-hypanthium (post anthesis, corolla and stigma absent); **F** fruit, side view; **G** fruit indumentum, detail; **H** pyrene, ventral view; **I** pyrene, side view; **J** pyrene plan view showing obscure lid and crest. All from *McPherson* 17011 (K). Drawn by Andrew Brown.

**Fig. 2.**
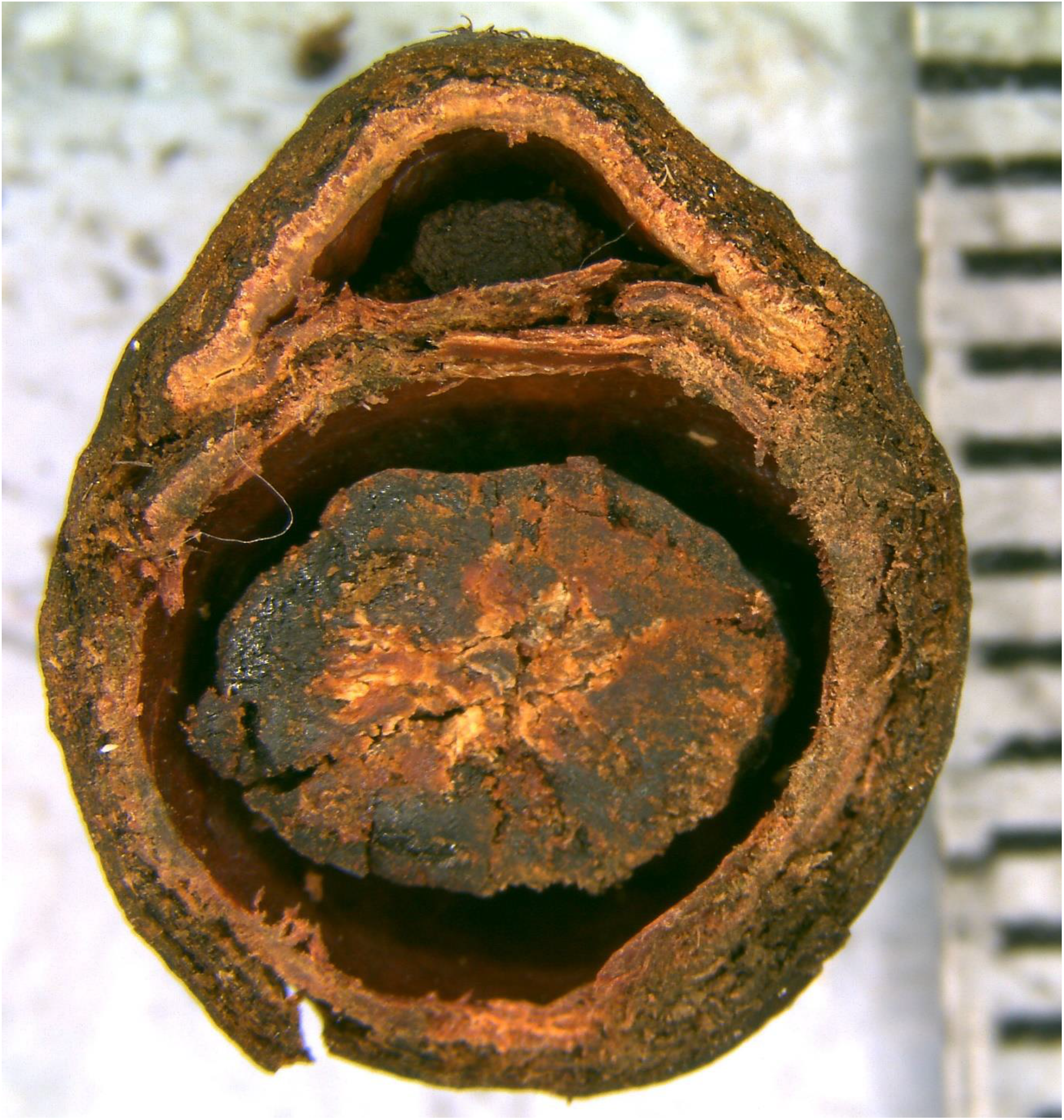
*Keetia gordonii* transverse section of 1-seeded fruit. Note the thick pyrene wall and the dense tanniniferous outer part of the seed endosperm enclosing a small white non-tanniniferous area with the embryo. Photo of *McPherson* 17011(K). Scale bar units = 1 mm.

##### RECOGNITION FEATURES

*Keetia gordonii* Cheek is unique among all known species of the genus for its massively thick, coriaceous leaf blades (vs papery in most other species, in this respect resembling *Psydrax* Gaertn.), and also for the globose, smooth pyrenes (vs ovoid, colliculose in other species), in which the lid crest is so vestigial that it is almost imperceptible (vs lid crest conspicuous in other species), with seeds in which the tanniferous area is completely solid in the outer half of the seed (not diffuse or in radial bands).

##### DISTRIBUTION & HABITAT

Western Gabon, lowland evergreen littoral terra firma forest, sometimes on sand, 0 − 250 m elev.

##### COLLECTIONS STUDIED

**GABON**, Ogooue-Maritime, 32 road-km N of Igotchi-Mouenda, Bakker forestry concession, 02° 41’ S, 010° 30’ E, 250 m, fr. 16 May 1997, *McPherson* 17011 (K holo., barcode K000593367!; isotypes LBV!, MO, UPS); ibid Rabi-E, SE of Pechoud Camp, “Lianescent shrub about 3 m high, several m long, young branches more hairy than older branches. Fruit containing 1 seed and several ants”, 01° 56.6’ S, 9° 53.1’ E, 0 m, fl. buds, 28 Oct. 1990, *van Nek* 137 (WAG 2 sheets, registration numbers WAG0094622 and WAG0094621; LBV!); Estuaire, Arboretum Raponda-Walker, Parcelle des Conservateurs. Forêt littorale humide de terre ferme, sur sol sableux. « Liane ligneuse de plusieurs mètres, à rameaux étalés. Tige pleine, cylindrique, à feuilles modifiées réduites et cordiformes. Rameaux étalés, souvent à ramifications divariquées. Feuilles vert foncé à nervures en creux dessus, vert clair à nervilles sombres très lâches dessous. Grandes stipules vertes. Rameaux et feuilles à poils brun-roux. Stérile. Fréquent », 00°34’13”N 009°19’01”E, 54 m, sterile, 18 Oct. 2020, *Lachenaud* 3029 with Nguema, Paradis, Bissiemou, Ikabanga, Nguimbit & Yombiyeni (BR, BRLU, LBV!, MO, WAG)

##### CONSERVATION STATUS

The type locality (*McPherson* 17011) for *Keetia gordonii* is just within the boundary of the Moukalaba-Doudou National Park in southwestern Gabon. Examination of the latest available satellite imagery at the collection site (Landsat/Copernicus dated October 2013, viewed March 2024) on Google Earth Pro shows intact forest, with no obvious threats, even though the point is only c. 1.3 km distant from a major road. The park is in an area with a low human population, but high numbers of large wild animals (Terada et al. 2021). Another rare species at this location, in fact endemic to the Moukalaba-Doudou National Park, is *Impatiens floretii* N. Hallé & A.M. Louis (Hallé & Louis 1989; Cheek 2022).

The second site for *Keetia gordonii* (*van Nek* 137) is in an onshore oil and gas production zone acquired in Feb. 2024 by the national Gabon Oil Company from Assala Energy (https://www.offshore-technology.com/news/gabon-to-buy-assala-energy/ accessed 7 April 2024). Here, in an area approximately 14 km N to S, and 4 km W to E equating to the Rabi Kounga II field, forest has been cleared in patches of approximately 120 m x 120 m, with the collection site about 100 m from one such clearing. However, this information is based on Landsat/Copernicus imagery dated April 2013 on Google Earth Pro (accessed 7 April 2024) so it is possible that in the intervening 11 years clearance of forest has become more extensive, possibly including the site at which *Keetia gordonii* was recorded in 1990.

At the third known location, the Forêt de la Mondah, (*Lachenaud* 3029), known since 2012 as the Raponda Walker Arboretum (Walters et al. 2016), habitat clearance and degradation has occurred due to its close proximity (c. 15 km) to the metropolis of Libreville which draws upon its trees for timber and firewood. Since created as a protected area in 1934, the forest has been reduced in size, losing 40% of its area in 80 years (Walters et al. 2016). These threats appear to be ongoing. The Raponda Walker forest is part of the Libreville region, which has the highest botanical specimen collection density in Gabon, with 5359 specimens recorded in digital format. It also has the highest level of diversity of both plant species overall and of endemics (Sosef et al. 2006). The coastal forests of the Libreville area are known to be especially rich in globally restricted species (Lachenaud et al. 2013). Despite this being the most intensively sampled part of Gabon, the species documented by Lachenaud et al. (2013) have also not been seen in some cases for several decades, or in the case of one species, since 1861, and are possibly extinct.

Given the ongoing threats identified above at two of the three locations, an area of occurrence of 12 km^2^ using the IUCN designated 4 km^2^ cells, we here assess the species provisionally as Endangered (EN B2a,b(i-iv)). The extent of occurrence is calculated as 6,964.3 km^2^ using Bachmann et al. (2011), exceeding the threshold for Endangered. It is possible that the species also qualifies as Endangered under criterion D on the basis of less than 50 mature individuals being recorded.

##### ETYMOLOGY

Named for the collector of the type specimen, Gordon McPherson an acclaimed botanist who collected herbarium specimens for Missouri Botanical Garden, not only in Gabon but also in countries such as Madagascar (see further notes below).

### Notes

*Keetia gordonii* does not conform to Bridson’s *Keetia* rule (Bridson 1986) in that the pyrene lid in this species, positioned on the top of the seed (and not the ventral face) is not correlated with presence of finely reticulate and conspicuous tertiary nerves. In fact, the nerves are very coarsely or not at all reticulate (Fig. 1A).

The new species remains incompletely known since flowering material is currently lacking. Although the notes of *van Nek* 137 talk of fruits, only immature inflorescences were present on the material examined. *Lachenaud* 3029 has a different aspect to the other two specimens but this may be due to it being at a different stage (sterile).

Gordon McPherson was a prolific collector for Missouri Botanic Garden where the top set of his specimens would be expected to be found. According to Missouri Botanic Garden’s Tropicos database (https://www.tropicos.org/collection/104802, accessed 29 March 2024), 8 duplicates were made, however, we have not been able to track down eight duplicates, but only three (see specimen citation above). Searches on the Naturalis bioportal (WAG https://www.naturalis.nl/en/science/bioportal), and the Meise (BR) specimen database (https://www.botanicalcollections.be/#/en/search/specimen) did not produce records of isotypes, contrary to expectations. Potentially there are five additional isotypes that have not been traced apart from the holotype at K, and isotypes recorded at MO and UPS. Alternatively, it may be that duplicates were destroyed, lost, or mislaid between being collected and distributed.

Gordon McPherson appears to have remained at the site of the type locality collecting specimens from 12 to19 May 1997 (https://vmtropicar-proto.ird.fr/gabon/collection/collecteurs/view/48?page=21 accessed March 2024) making specimens in his number range 16959 to 17019

## Discussion

The taxonomic placement of *Keetia gordonii* is probably with what is here designated as the *Keetia hispida* group, including the *K. hispida s*.*l*. complex itself, *K. bakossiorum* Cheek, *K. ornata* Bridson & Robbr. and two other currently unrecognized species mainly in Cameroon, which together share long, densely, often wiry-hairy, stems, large leaf blades, and very large fruits in which rather than the apex of the 2-seeded fruits being retuse or indented as usual in the genus (Bridson 1986), it appears shortly rostrate, due to the apex being prolonged by a persistent and accrescent, long calyx tube (Fig. 1F). However, these similarities may be the result of convergence rather than shared recent ancestry. Densely sampled species-level molecular phylogenetic studies of the genus are desirable.

## Conclusion

It is important to identify and formally name species so that IUCN extinction risk assessments can be made and accepted, allowing the possibility of resources being allocated for their conservation if they are threatened Cheek et al. (2020). About two thirds of new plant species being described currently are threatened with extinction at the point of publication (Brown et al. 2023). Species of plant unique to Gabon are already seemingly globally extinct e.g. *Pseudohydrosme buettneri* Engl. and *P. bogneri* Cheek & Moxon-Holt (Cheek et al. 2021; Moxon-Holt & Cheek 2021) although there is always the hope that they will be rediscovered. Completion of formal naming of species in Gabon is vital if further extinctions are to be avoided and if Gabon is not to follow its neighbour, Cameroon, which has the highest number of global extinctions in continental tropical Africa (Humphreys et al. 2019) and where new global plant species extinctions continue to be recorded (Cheek et al. 2017; 2018b; 2018c; 2019; Murphy et al. 2023).

## Acknowledgements

We thank Shigeo Yasuda for preparing pyrenes for study, and Hannah Bevan for calculating the extent of occurrence. Sebastian Hatt helped prepare Figure 2.

The authors declare that they have no conflict of interest.

